# Fine-scale cellular deconvolution via generalized maximum entropy on canonical correlation features

**DOI:** 10.1101/2024.06.07.598010

**Authors:** Jack Kamm

## Abstract

We propose a method for estimating probability distributions over single cells, which we apply to fine-scale cellular deconvolution, which quantifies the composition of external bulk RNAseq samples at high resolution (i.e. at the single-cell or neighborhood level). Our method is based on a computationally-efficient convex optimization problem, which is also generalization of the Maximum Entropy method. Our method has a much higher resolution than traditional approaches that require computing gene expression profiles at the cell-type level, and also compares favorably to recent high-resolution cellular deconvolution methods, with orders-of-magnitude speedup in computational efficiency. We implement this method in a Python package quipcell, available at https://github.com/genentech/quipcell.

## 1 Introduction

Single-cell RNAseq (scRNAseq) is a powerful tool for understanding heterogeneous biological samples consisting of diverse tissues, cell types, or cell states. A common application of scRNAseq data is to quantify differences in cell composition between samples or conditions. Two variants of this task are *differential abundance* analysis and *cellular deconvolution* analysis (also known as *bulk deconvolution*). In differential abundance analysis, we seek to compare cell composition between samples within the same single-cell dataset (Amezquita et al. (2023), Chapter 6). By contrast, in bulk deconvolution analysis, we use single-cell data to estimate the composition of external bulk RNAseq samples (Maden et al., 2023; Nguyen et al., 2024).

One straightforward approach to estimating sample composition, in both differential abundance and bulk deconvolution, is to bin cells into discrete cell types or clusters, and then to quantify the abundance of these discrete groups across samples. However, this coarse-grained approach can miss fine-scale differences in cell type composition, such as from continuous cell differentiation processes.

In recent years, several high-resolution methods have been developed for differential abundance. Typically, these methods involve building a nearest-neighbor graph of the single cells, and then quantifying the abundance of samples along the graph. In particular, the methods MELD (Burkhardt et al., 2021) and CNA (Reshef et al., 2022) use a random walk on the nearest neighbor graph to represent each sample as a probability distribution over cells, while miloR (Dann et al., 2022) breaks up the graph into overlapping neighborhoods and measures the sample frequency and spatial correlation of the neighborhoods. DA-seq (Zhao et al., 2021) is another high-resolution differential abundance method that makes use of nearest-neighbor information, by fitting logistic regression models in neighborhoods around each cell to distinguish between samples.

By contrast, there are relatively few bulk deconvolution methods that work at single cell resolution. Instead, most deconvolution methods aggregate single cell data into expression profiles of discrete cell types, and then model the bulk samples as mixtures of these discrete profiles (Altboum et al., 2014; Newman et al., 2019; Wang et al., 2019; Jew et al., 2020; Dong et al., 2021; Erdmann-Pham et al., 2021). One notable exception is CPM (Frishberg et al., 2019), which uses Support Vector Regression to model the bulk samples as a linear combination of reference single cells. However, CPM was designed to distinguish continuous cell states within cell types, and may not work well on datasets containing very diverse cell types. More recent high-resolution deconvolution methods include ConDecon (Aubin et al., 2023), which learns a map from rank-correlation scores to a probability distribution over single cells; and MeDuSa (Song et al., 2023), which deconvolves bulk samples along a linear continuous pseudotime trajectory using a mixed effects model. In addition, spatial deconvolution methods, such as Tangram (Biancalani et al., 2021), bear some similarities to high-resolution bulk deconvolution, but operate in a different regime with distinct challenges (many voxels with few cells per voxel) as well as advantages (additional spatial information, well-matched single-cell and spatial samples).

We propose a novel method for bulk deconvolution, quipcell, that is a convex optimization problem and a generalization of the Maximum Entropy method (Jaynes, 1957). Quipcell represents each sample as a probability distribution over some reference single-cell dataset, similar to the MELD differential abundance method (Burkhardt et al., 2021), or the ConDecon bulk deconvolution method (Aubin et al., 2023). However, quipcell fits this distribution using an alternative approach that only relies on solving an efficient convex optimization problem, instead of constructing nearest neighbor graphs or fitting predictive machine learning models.

Quipcell is based on the Maximum Entropy method (MaxEnt), which fits the “least informative” probability distribution consistent with observed data, and has been widely applied in areas ranging from statistical physics (Jaynes, 1957) to ecology (Harte and Newman, 2014). In particular, MaxEnt maximizes the Shannon entropy *H* (Shannon, 1948) as the metric of uninformativeness. However, other types of entropy can also be used to measure information, such as the Rényi entropy *H*_*α*_ (Rényi, 1961), a generalization of the Shannon entropy that is closely related to *α*-diversity metrics commonly used in ecology and metagenomics (Jost, 2006). Quipcell performs bulk deconvolution by finding the probability distribution over single cells that maximizes the Rényi entropy *H*_*α*_ (with the special case *α* = 1 being equivalent to standard MaxEnt). This approach can also be formulated as a special case of the Generalized Cross Entropy method for density estimation (Botev and Kroese, 2011), which involves matching distributional moments while minimizing an *f* -divergence (Csiszár, 1964) from the uniform distribution.

A key aspect of this density estimation procedure is the embedding space used to represent the single cells. Quipcell requires this embedding to be a linear transformation of the original single cell data. We show that an effective approach is to use Canonical Correlation Analysis (CCA) (Hotelling, 1936) to find directions in gene expression space that are highly correlated with celltype-relevant features. In the case that the single cell data is annotated with discrete celltype labels, we perform CCA between the gene expression matrix and the indicator matrix of the celltype labels; this is also equivalent to the multiclass Linear Discriminant Analysis (LDA) (Rao, 1948). In the case where the single cell data does not have discrete celltype labels, but is instead annotated with a continuous “pseudotime”, we perform CCA between the gene expression matrix and a cubic B-spline basis (De Boor, 1978) to extract the embedding space.

For our application (Section 3), we consider three datasets. First, we apply quipcell to the Human Lung Cell Atlas dataset (Sikkema et al., 2023), which is a highly heterogeneous reference set consisting of many studies and technologies. We show that our method has good accuracy and runtime properties on this heterogeneous reference, especially compared to existing high-resolution bulk deconvolution methods. Next, we apply quipcell to the problem of pseudotime deconvolution on datasets of stem cells (Chu et al., 2016) and monocytes (Oetjen et al., 2018), and show comparable performance to the pseudotime deconvolution method MeDUSa (Song et al., 2023).

Quipcell is implemented as a Python package, available at https://github.com/genentech/quipcell.

## 2 Methods

Let the single-cell reference set **X** ∈ ℝ^*n*×*p*^ be a matrix of *n* cells by *p* features. For example, **X** could be normalized scRNAseq gene counts, in which case *p* would equal the number of genes.

Let ***µ***_*s*_ ∈ ℝ^*p*^ be the bulk expression of the sample *s* on these *p* features. We aim to represent *s* as a probability distribution ℙ_*s*_ over ℝ^*p*^ such that 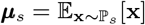. Intuitively, ℙ_*s*_ represents a probability distribution over single cells, and the bulk expression levels ***µ***_*s*_ are measured as an average over the single cells in *s*. However, note that ℙ_*s*_(**x**) is *cell-size biased*, because larger cells contribute more reads to the bulk expression ***µ***_*s*_. In particular, ℙ_*s*_ represents the probability that a random *read* from *s* is sampled from a cell at **x**, rather than the probability that a random *cell* from *s* is sampled at **x**. In Section 2.3, we discuss how to correct the cell-size bias and convert the read-level probabilities to cell-level probabilities.

We approximate ℙ_*s*_ by a distribution 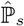 with support on the *n* single cells (row vectors) of the reference set **X**. We let **ŵ**_*s*_ = (*ŵ*_*s*,*c*_)_1≤*c*≤*n*_ ∈ ℝ^*n*^ denote the corresponding vector of probability weights 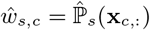, where **x**_*c*,:_ denotes the *c*-th row vector (cell) in **X**. Intuitively, we interpret *ŵ*_*s*,*c*_ as the probability that a random read from *s* originates from a small neighborhood around cell *c*. Furthermore, we use *ŵ*_*s*,*c*_ as importance sampling weights to approximate expectations over 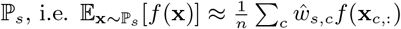.

We fit **ŵ**_*s*_ ∈ ℝ^*n*^ via the Generalized Cross-Entropy method for density estimation (Botev and Kroese, 2011); more specifically, we solve the convex optimization problem (for any fixed *ϵ* ≥ 0 and *α >* 0):

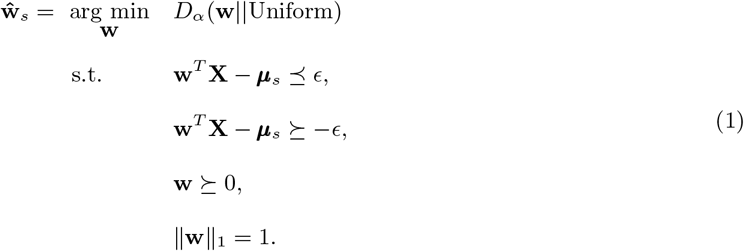

where *Dα*(**w**||Uniform) denotes the *α*-divergence of **w** from the discrete uniform distribution, and is defined as:

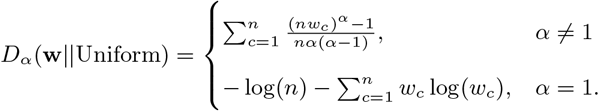

When *α* = 1, this is equivalent to the KL-Divergence and also (up to constant) the negative Shannon entropy −*H*(**w**), i.e. *D*_1_(**w**||Uniform) = *D*_*KL*_(**w**||Uniform) = −*H*(**w**) + *H*(Uniform).

Note that for fixed *α, D α* (**w**||Uniform) monotonically increases with the negative Rényi entropy 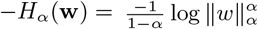 (Rényi, 1961), so we could equivalently use −*H*_*α*_(**w**) as the objective function instead of *D α* (**w**||Uniform). Therefore, quipcell can viewed as a “Maximum Rényi Entropy” method. Intuitively, quipcell searches for the least informative **w** consistent with the moments ***µ***_*s*_, where “informativeness” is defined as the negative Rényi entropy −*H α* (**w**), or equivalently as the *α*-divergence from the Uniform distribution *D α* (**w**||Uniform).

Two notable special cases of *Dα* (·||·) are the KL-divergence (*α* = 1), in which case (1) is the Maximum Entropy method, and Pearson’s *χ*^2^-divergence (*α* = 2), in which case (1) is a Quadratic Program (QP) and can be solved especially efficiently. Botev and Kroese (2011) particularly favor using *α* = 2, and point out several advantages over other *f* -divergences (including the KL-divergence), such as computational efficiency, robustness to outliers, and the fact that minimizing this objective also guarantees an upper bound on both the KL-divergence and the *L*_1_ (Total Variation) distance.

For the default values of the parameters *α, ϵ* in (1), quipcell uses *α* = 2 (following Botev and Kroese (2011)) and ϵ = 0 (requiring exact equality for the monent constraint). We use these default settings for the examples in the remainder of this manuscript unless stated otherwise. In this case, (1) can be rewritten as the quadratic program

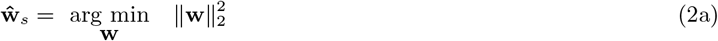

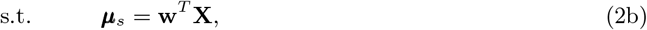

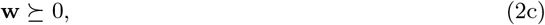

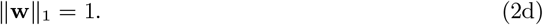

### 2.1 Solution via convex optimization

*D*_*α*_ (**w**||Uniform) is convex in **w**, so (1) can be efficiently solved via convex optimization, and model diagnostics can also be obtained by inspection of the Lagrangian dual solution.

Quipcell uses CVXPY (Diamond and Boyd, 2016) to solve (1) and fit **ŵ**_*s*_. The default case *α* = 2 is a Quadratic Program (QP) and can be solved particularly efficiently; CVXPY uses the OSQP solver (Stellato et al., 2020) by default in this case. In addition, the case *α* = 1 (Maximum Entropy) can be solved efficiently and directly via its Lagrangian dual (Jaynes (1957); see also Section 5.5.5 of Boyd and Vandenberghe (2004)). For this case, quipcell uses scipy (Virtanen et al., 2020) and JAX (Bradbury et al., 2018) to run L-BFGS-B (Byrd et al., 1995) on the dual. For general *α* ∉ {1, 2}, quipcell uses CVXPY with the ECOS solver (Domahidi et al., 2013) by default; however, quipcell also allows users to specify alternative solvers and options to CVXPY for optimization.

Any solution to (1) is guaranteed to be unique, due to the strict convexity of *Dα* (**w**||Uniform). The strict convexity can be seen by noting that the first derivative is strictly increasing along any direction **z**: defining 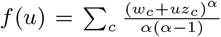, its second derivative is 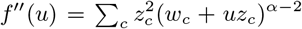, which is strictly positive on {*u* : **w** + *u***z** ⪰ 0} Lebesgue almost everywhere.

The stability of the solution can be analyzed via the Lagrangian dual of (1). Locally, the dual solution λ^*^ gives the gradient of the optimum with respect to the constraints (see Section 5.6.3 of Boyd and Vandenberghe (2004)). In particular, if a dual value corresponding to the *i*th coordinate of ***µ***_*s*_ is large, then small perturbations of the *i*th feature may cause large changes in the result. quipcell provides functionality for users to inspect the dual solution λ^*^ to aid interpretation.

### 2.2 Linear transformations and canonical correlation features

In the prior exposition, we assumed that **X** and ***µ***_*s*_ represented normalized counts of *p* genes. In practice *p* can be large, and it may be desirable to use a lower dimensional representation to improve speed and prevent overfitting.

By linearity of expectation, we may replace **X** and ***µ***_*s*_ with any linear transformation **X***A*^*T*^ and *A****µ***_*s*_, because 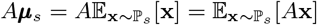.

In our applications below, we set *A* = *A*_2_*A*_1_, where *A*_1_ are PCA gene-loadings, so that 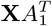 gives the loadings of each cell along the principal components, and *A*_2_ are loadings of a CCA between the PCA cell-loadings 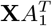 and some informative set of cell annotations *B* ∈ ℝ^*n*×*k*^. In our first application (Section 3.1), we set *B* to be the indicator (one-hot) matrix of the celltype annotations from the Human Lung Cell Atlas. Note that the CCA of an indicator matrix is equivalent to the multiclass Linear Discriminant Analysis (Bach and Jordan, 2005). In our second example (Section 3.2), we set *B* to be a cubic B-spline representation of the cell’s pseudotime coordinate.

Note that quipcell only works on linear transformations of the normalized gene counts. In particular, the gene counts should **not** be log-transformed after normalization.

### 2.3 Correcting cell-size bias to estimate cell-level probabilities

As noted earlier, the weights *ŵ*_*s*,*c*_ represent the probability of sampling a random read from *s* at cell *c*. To see this, consider the case where *s* is comprised of a subset of cells from **X**, i.e. *s* ⊆ {1,, …, *n*}. Let **Y** be the raw single cell gene counts, and *n*_*c*_ = ∑ _*j*_ *Y*_*c*,*j*_ the size factor for cell *c*. Let 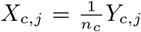 be the normalized gene counts. Then we have that 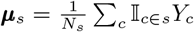, where *N*_*s*_ = ∑ _*c∈s*_ *n*_*c*_. If we then consider the optimization problem (1) with ***µ***_*s*_ and **X**, a feasible point is 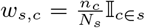, which is the probability that a random read of *s* comes from cell *c*. Therefore we interpret the inferred weights *ŵ*_*s*,*c*_ as read-level rather than cell-level probabilities. The same argument holds if we replace **Y, X, *µ***_*s*_ with any linear transformation **Y***A*^*T*^, **X***A*^*T*^, *A****µ***_*s*_.

Since the weights *ŵ*_*s*,*c*_ correspond to the probability of sampling reads rather than cells, they should only be used to compare relative abundances *between* samples, and cannot directly compare cell-type abundances *within* a single sample.

To enable within-sample comparisons, we can also obtain cell-level weights 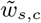 by renormalizing the read-level weights *ŵ*_*s*,*c*_. In particular, if we have cell-specific size factors *m*_*c*_, we can convert the *read* -level abundances *ŵ*_*s*,*c*_ to *cell* -level abundances 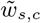 by

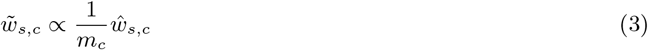

where the size factor *m*_*c*_ is proportional to the number of UMIs coming from cell *c*.

Naively, we could simply set *m*_*c*_ to be the number of UMIs from cell *c*; however, this can be problematic for heterogeneous reference sets, such as the Human Lung Cell Atlas (HLCA). For example, Figure A.2 shows that the different studies within the HLCA have very different sequencing depths and cell-type distributions.

We therefore use a Poisson GLM to estimate cell-specific size factors *m*_*c*_ that account for different sequencing depths between batches or samples. In particular, we estimate the number of UMIs *n*_*c*_ as a function of the cell’s sample *S*_*c*_ and its low-dimensional representation *XA*^*T*^ :

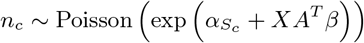

where we estimate the sample-specific normalization terms *α*_*S*_ and the feature coefficients *β*. We then set the cell-specific size factor *m*_*c*_, after controlling for the sample sequencing depth, as

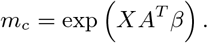

We then use these cell-specific size factors *m*_*c*_ to convert between read-level and cell-level abundances with equation (3).

Note that the renormalized cell-level weights may be less accurate than the read-level weights – see Figure 2 for a comparison, and Maden et al. (2023) for a discussion of challenges on converting read-level to cell-level abundances. See also Maden et al. (2024) for a recent method to correct cell-specific size factors in traditional (coarse-grained) bulk deconvolution.

## 3 Results

### 3.1 Application to Human Lung Cell Atlas

We applied quipcell to using the Human Lung Cell Atlas (HLCA) (Sikkema et al., 2023) as the reference single cell data set. It originally comprised 580K cells from 166 samples (107 donors) across 11 studies, but we held out the `Krasnow_2020` study (Travaglini et al., 2020) from the dataset as a validation set for bulk deconvolution, resulting in a reference set with 520K cells. For the cell-feature matrix **X**, we used the loadings from the top 15 components of a multiclass Linear Discriminant Analysis (LDA), using the cell types from Sikkema et al. (2023) (shown in Figure 1a) as the class labels. The held-out `Krasnow_2020` study (Travaglini et al., 2020) consisted of 5 samples, from which we generated pseudobulk samples to benchmark bulk deconvolution with a known ground truth. See Appendix A.1 for more details about processing steps, and Figure 1a for a UMAP of the HLCA dataset.

**Figure 1:**
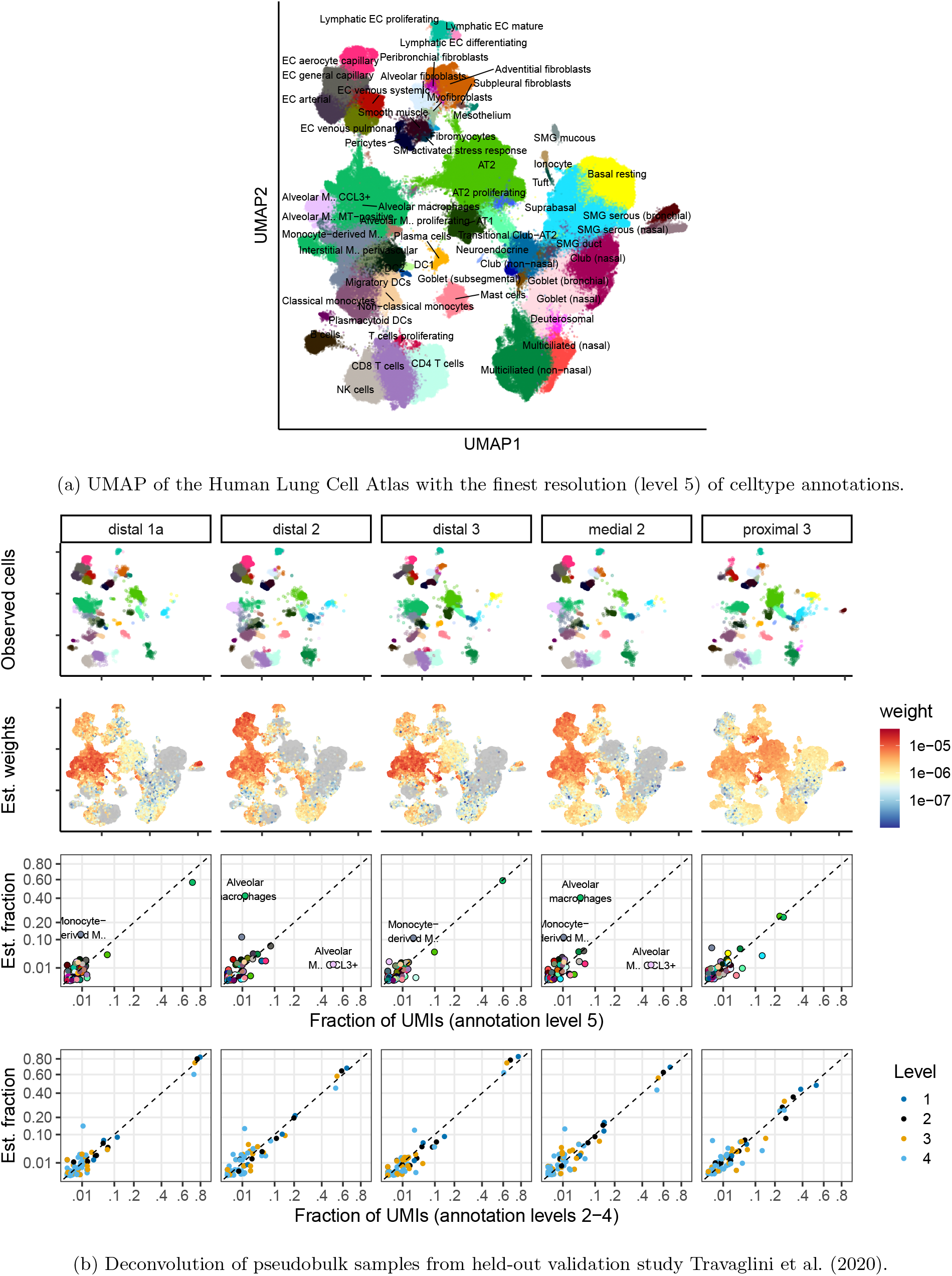
Results from deconvolving the held-out validation study Travaglini et al. (2020) on the rest of the Human Lung Cell Atlas (Sikkema et al., 2023). (1a) UMAP and cell type labels downloaded from the original publication. (1b) Top row shows the cells from each of the 5 held-out samples. 2nd row shows estimated weights *ŵ*_*s*,*c*_ for each sample; weights below 1e-8 were set to 0 and are colored gray. 3rd row shows the fraction of UMIs coming from each cell type (annotation level 5), vs the estimated fraction based on the weights *ŵ*_*s*,*c*_. The x and y axes are square-root-scaled. Note quipcell has trouble distinguishing some macrophage subtypes, especially the “Alveolar macrophages” and “Alveolar Macrophage CCL3+” subtypes (see Appendix A.2). 4th row shows the accuracy of bulk deconvolution results at coarser resolutions of cell-type annotation (annotation levels 2-4).

**Figure 2:**
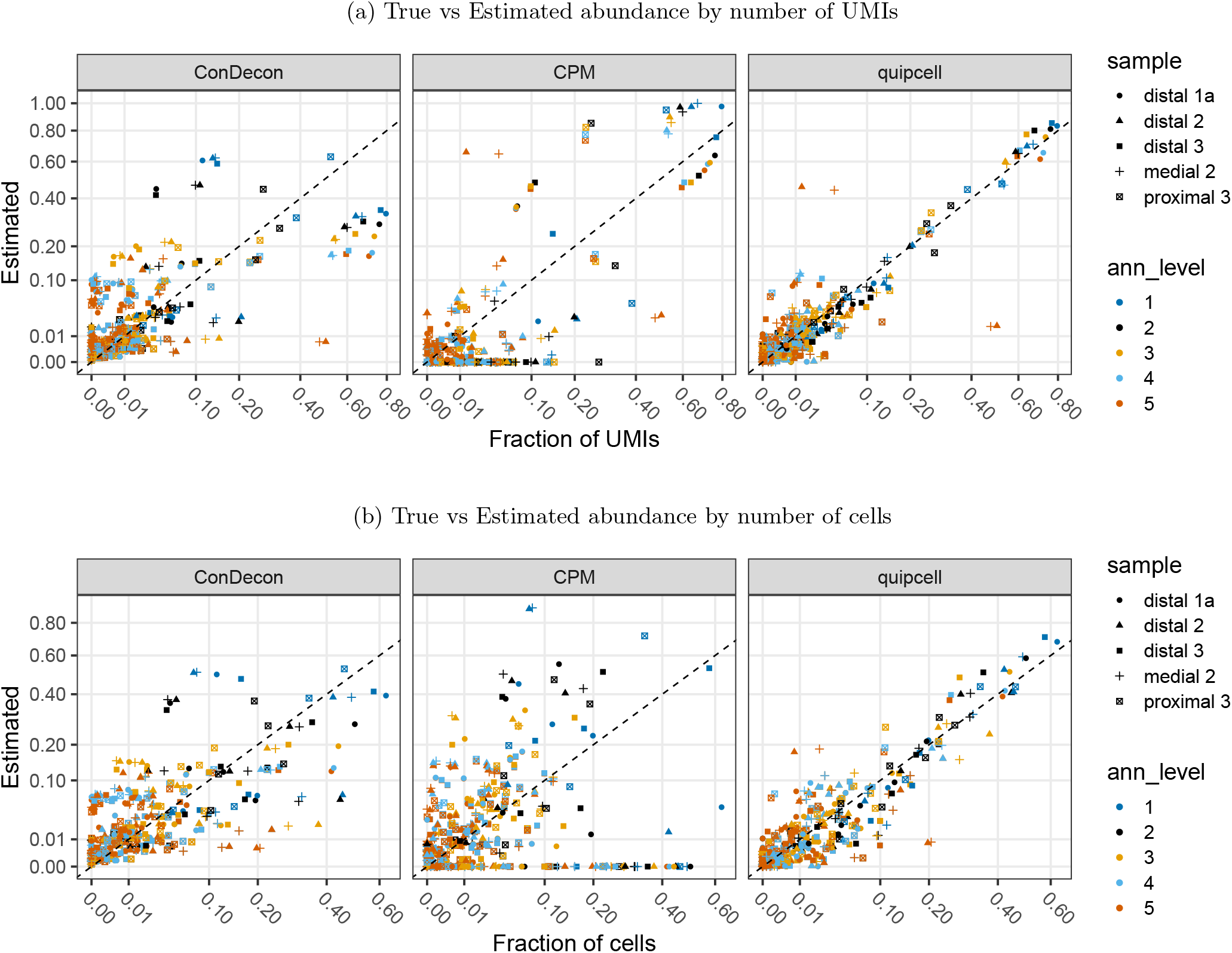
Comparison of our density estimation method with CPM (Frishberg et al., 2019) and ConDecon (Aubin et al., 2023). For each sample in the held-out validation study (Travaglini et al., 2020), we compare the estimated fraction of UMIs (top) or cells (bottom) coming from each celltype across several annotation levels (higher level means finer resolution). *x* and *y* axes are transformed by a square-root scaling.

We show the results of the deconvolution in Figure 1b. Overall, the estimated abundances are reasonable, though we note that the quipcell estimates become more accurate (closer to the diagonal) as the resolution becomes coarser (with annotation level 1 being the coarsest level of annotation, and annotation level 5 being the finest; see also Figure A.3). In particular, the method struggles to distinguish some macrophage subpopulations at the finest annotation resolutions, most notably “Alveolar macrophage CCL3+” vs. other Alveolar macrophages, and to a lesser extent “Interstitial macrophages” (more specifically “Monocyte-derived macrophages”) from other macrophages. Part of the trouble stems from the fact that “Alveolar macrophage CCL3+” cells mainly appear in the validation set, and are relatively rare in the reference set (Table A.2), which causes none of the top 15 LDA components to separate these populations well (Figure A.5).

In Appendix A.3, we illustrate how examining the dual variables can be used to inspect model diagnostics in this example, and in Appendix A.4, we compare how the results change when using different *α*-divergences.

#### 3.1.1 Comparison with existing fine-resolution deconvolution methods

We compare our method with two existing fine-resolution deconvolution methods: CPM (Frishberg et al., 2019) and ConDecon (Aubin et al., 2023). We plot the accuracy of these methods in Figure 2, which shows our method to more accurately estimate abundances in general – in particular, notice that quipcell has points closer to the diagonal. Note that we ran all methods on a random 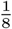 of the reference set for this comparison, so that all methods could finish running with a reasonable amount of time and memory.

CPM (Frishberg et al., 2019) solves a similar QP as quipcell, but uses the gene counts directly (instead of a feature space like LDA), and also lacks the constraints (2c), (2d). This means the CPM-inferred weights don’t sum to 1, and generally include negative weights. To compare cell-type proportion estimates, we summed the CPM weights by cell-type, set the negative cell-types to 0, and renormalized the positive cell-types to sum to 1. Note that setting celltypes with negative probability to 0 may be the culprit for the bottom horizontal stripe in the CPM panels of Figure 2. Note also that CPM does make use of a lower-dimensional embedding of the reference data, but only to subsample the reference single-cells according to a grid, and this embedding is required to be 1- or 2-dimensional. We used the original UMAP from the HLCA (Figure 1a) for this 2-dimensional embedding when running CPM.

ConDecon (Aubin et al., 2023) is a more recent high-resolution deconvolution method, that learns a map from the rank correlations between bulk and single-cell samples to a probability distribution on single cells. The method is trained on random resamplings of the single-cell training set. It also requires an embedding space of the single-cell data; we used the same 15-dimensional LDA embedding for this, as in quipcell. We ran ConDecon with default parameters.

**Table 1:**
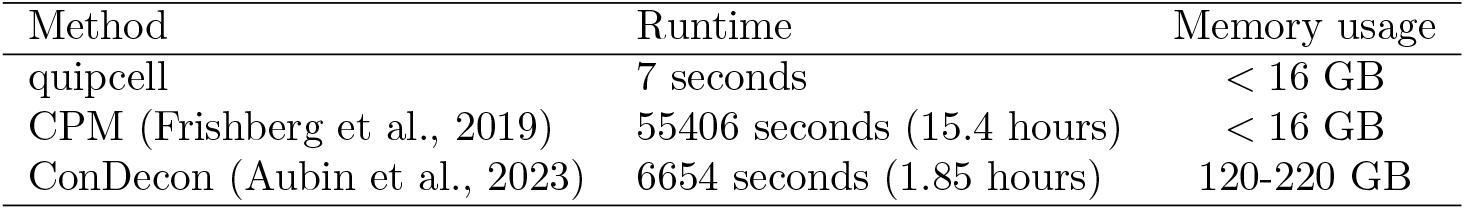
Comparison of runtime and memory usage to perform deconvolution on the 5 validation samples. Memory usage is approximate: quipcell and CPM were run in jobs that were allocated 16 GB memory, but in fact use much less memory. ConDecon failed when run in a job allocated 120 GB memory, but succeeded when the memory limit was increased to 220 GB. Note that all methods were run on the same random 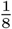 subset of the reference for this comparison.

Note that the QP of quipcell and CPM estimate UMI-level distributions, while ConDecon estimates cell-level distributions. However, the cell-level and UMI-level representations can be converted between each other by using cell-specific size factors (Section 2.3). We use the same cell size factors, estimated by the Poisson GLM of Section 2.3, for all 3 methods. In Figure 2, we show the accuracy of the three methods on both UMI-level and cell-level abundances.

In terms of runtime, our QP-based method was more performant than either CPM or ConDecon, and ran in about 7 seconds for 5 bulk samples on a job allocated 16GB RAM (about the same as a laptop). By contrast, CPM took 15 hours to run, likely due to its lack of sparse matrix support. ConDecon took about 110 minutes to run, and also required significant memory usage (between 120-220GB RAM).

### 3.2 Application to pseudotime deconvolution

We next applied quipcell to the problem of deconvolving a bulk sample along a 1-dimensional pseudotime. We compared quipcell against the recent pseudotime deconvolution method MeDuSA (Song et al., 2023), on two datasets featured in the MeDuSA tutorial vignettes: a dataset of human stem cells (Chu et al., 2016), and a dataset of monocytes from human bone marrow (Oetjen et al., 2018). Both datasets contained matched single-cell and bulk samples, and we applied deconvolution to the bulk RNAseq samples, taking the matched single-cell samples as the ground truth.

In both cases, we used the pseudotime annotations provided with the MeDuSA vignette, and ran MeDuSA with the same settings as in the vignette (`markerGene=NULL`, `span=0.35`, `resolution=50`, `smooth=TRUE`, `fractional=TRUE`).

To run quipcell, we used canonical correlation features derived from a CCA between the gene-expression matrix and a cubic B-spline basis of the pseudotime coordinate (computed with the R `fda` package, Ramsay and Silverman (2005)), and report the cell-size corrected weights (Section 2.3). In both cases, we used a basis of 16 splines and selected the top 3 CCA components (using more CCA components resulted in overly sparse weights).

Figure 3 shows the deconvolution results on the monocyte dataset, while Figure 4 shows the deconvolution results on the stem cell dataset. To compare quipcell and MeDuSA, we used the pseudotime bins output by MeDuSA, and computed the total quipcell weights within each bin, as well as the fraction of ground truth cells. We computed the summed absolute (*L*_1_) and squared (*L*_2_) errors of the predicted fractions against the ground truth fractions for both methods. In the monocyte dataset (Figure 3), quipcell generally had lower errors than MeDuSA, while in the stem cell dataset (Figure 4) the results were mixed, with MeDuSA being more accurate on some samples, and less accurate on others.

**Figure 3:**
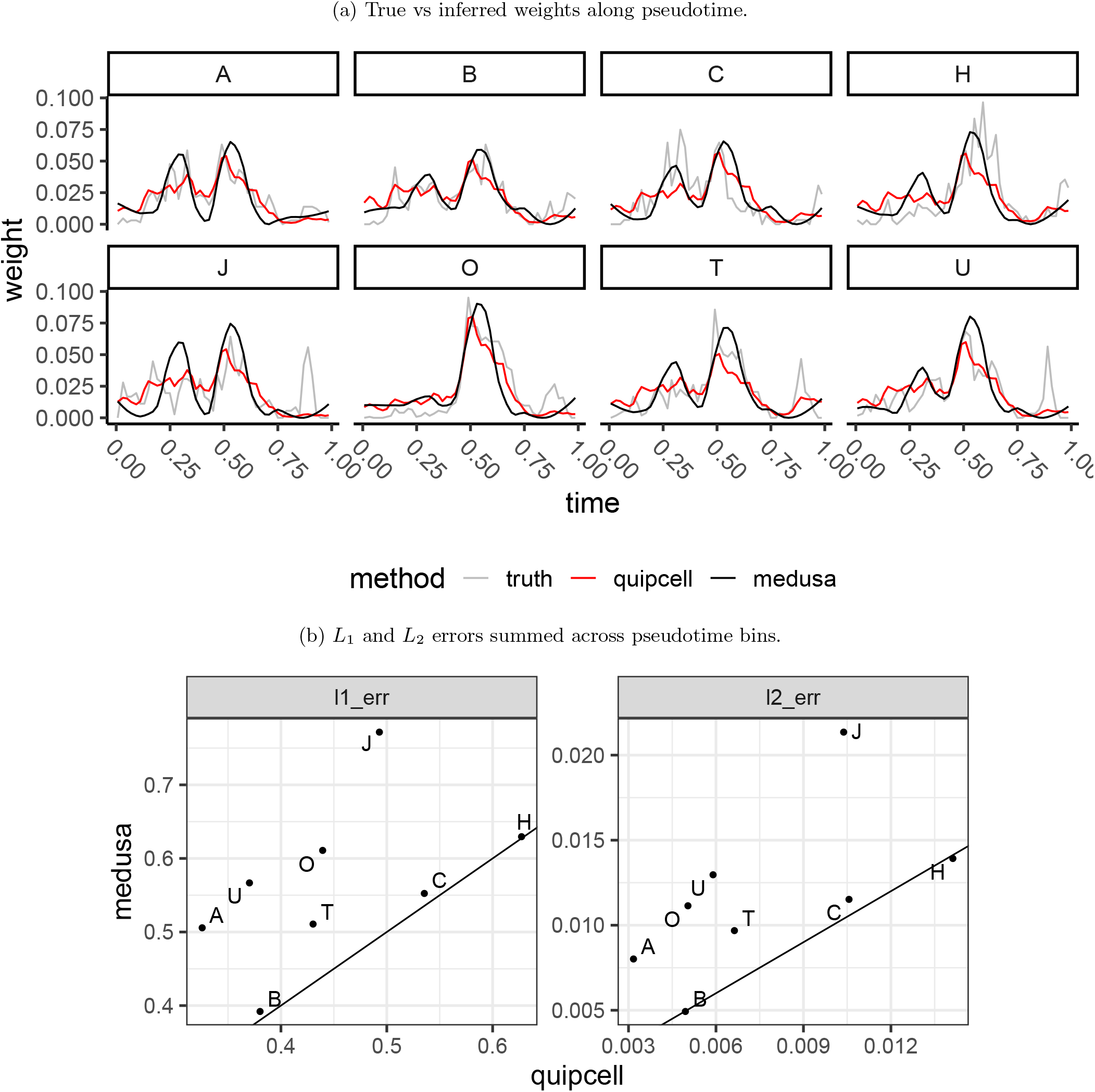
Comparison of quipcell and MeDuSA on the 8 bulk samples from the monocyte dataset.

**Figure 4:**
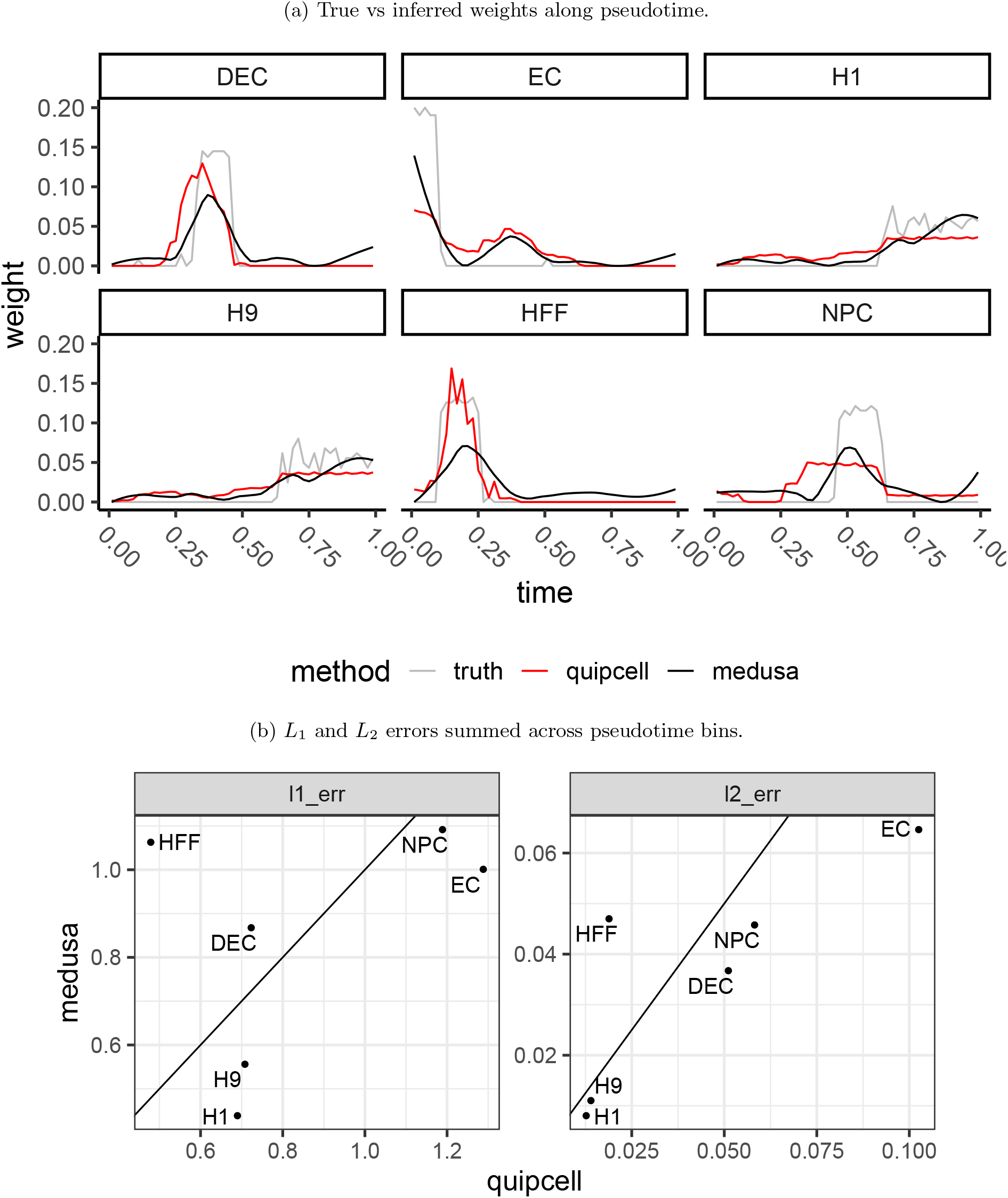
Comparison of quipcell and MeDuSA on the 6 bulk samples from the stem cell dataset.

## 4 Conclusions

We present quipcell, a distribution-fitting method for single-cell data based on moment-matching and convex optimization, which can be applied for bulk deconvolution analyses. When combined with Canonical Correlation Analysis (or its special case, Linear Discriminant Analysis), our method performs well on a range of datasets, including highly heterogeneous single cell atlases, and compares favorably to other high-resolution bulk deconvolution methods in both computational efficiency and predictive accuracy.

One direction for future work would be to apply the generalized entropy framework to the problem of differential abundance, i.e. estimating density changes *within* a single-cell dataset. While many existing fine-scale differential abundance methods exist (Burkhardt et al., 2021; Reshef et al., 2022; Dann et al., 2022; Zhao et al., 2021), they typically involve quantifying abundances along a nearest-neighbor graph; a generalized entropy approach would represent an alternative, graph-free approach based on information-theoretic principles. Unlike in the bulk deconvolution case, we would not be limited to linear transformations of the gene counts **X**, since we could explicitly compute the expectation 𝔼_**x**∼𝕡 *s*_ [*f* (**x**)] even for nonlinear *f* . A notable special case would be to use a Reproducing Kernel Hilbert Space as the feature space – then the results of Botev and Kroese (2011) show that the solution is a *continuous* density estimator based on a Gaussian mixture model, which is also equivalent to a sparse weighted kernel density estimator with data-driven bandwidth. If this kernel is furthermore a graph diffusion kernel, the estimate may become related to random-walk-based differential abundance methods such as CNA (Reshef et al., 2022) and MELD (Burkhardt et al., 2021).

## Acknowledgments

I thank Bill Forrest, Oleg Mayba, and Jayaram Kancherla for their invaluable feedback.

## A Appendix

### A.1 Processing steps for HLCA dataset

Here we describe the processing steps used for our analyses of the HLCA dataset.

First, we used scanpy (Wolf et al., 2018) to normalize each cell by its total UMI count, select highly variable genes (using study as the batch variable, and otherwise default parameters; 2176 genes selected) and perform PCA with 100 PCs. Note that we did not apply log-scaling of the data; therefore, the PCA is a linear transformation of the normalized counts.

Next, we split the dataset into two cohorts:

1. A hold-out “validation” cohort consisting of samples from the `Krasnow_2020` study (Travaglini et al., 2020), used to generate pseudobulk samples ***µ***_*s*_ for deconvolution. We chose to hold out this study because it contained the fewest samples (5) while still having a diversity of cell types, making it easier to visualize the full results as in Figure 1b.
2. A “reference” cohort consisting of the remaining cells. The weights *ŵ*_*s*,*c*_ are computed on cells *c* coming from this reference cohort.
  a. For speed reasons, some analyses (Figure 2, Figure A.6), use a smaller reference consisting of a random 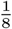 of the full reference.

We list the number of cells, samples, and studies in each subset in Table A.1.

**Table A.1:**
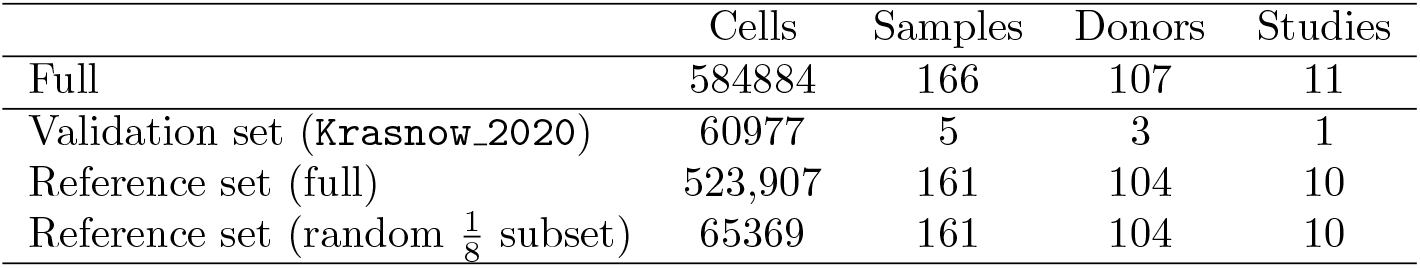
Table of number of cells, samples, and studies in each of the data subsets. The validation set consists of one held-out study, and is used to benchmark bulk deconvolution. For some analyses (Figure 2, Figure A.6), we only use a random 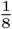 of the reference set to improve speed.

After excluding the hold-out validation cohort, we performed multiclass Linear Discriminant Analysis on the 100 PCs, using the HLCA cell types as the class labels. After examining a plot of the variances along each LDA component, we kept the top 15 components of the LDA (Figure A.1).

**Figure A.1:**
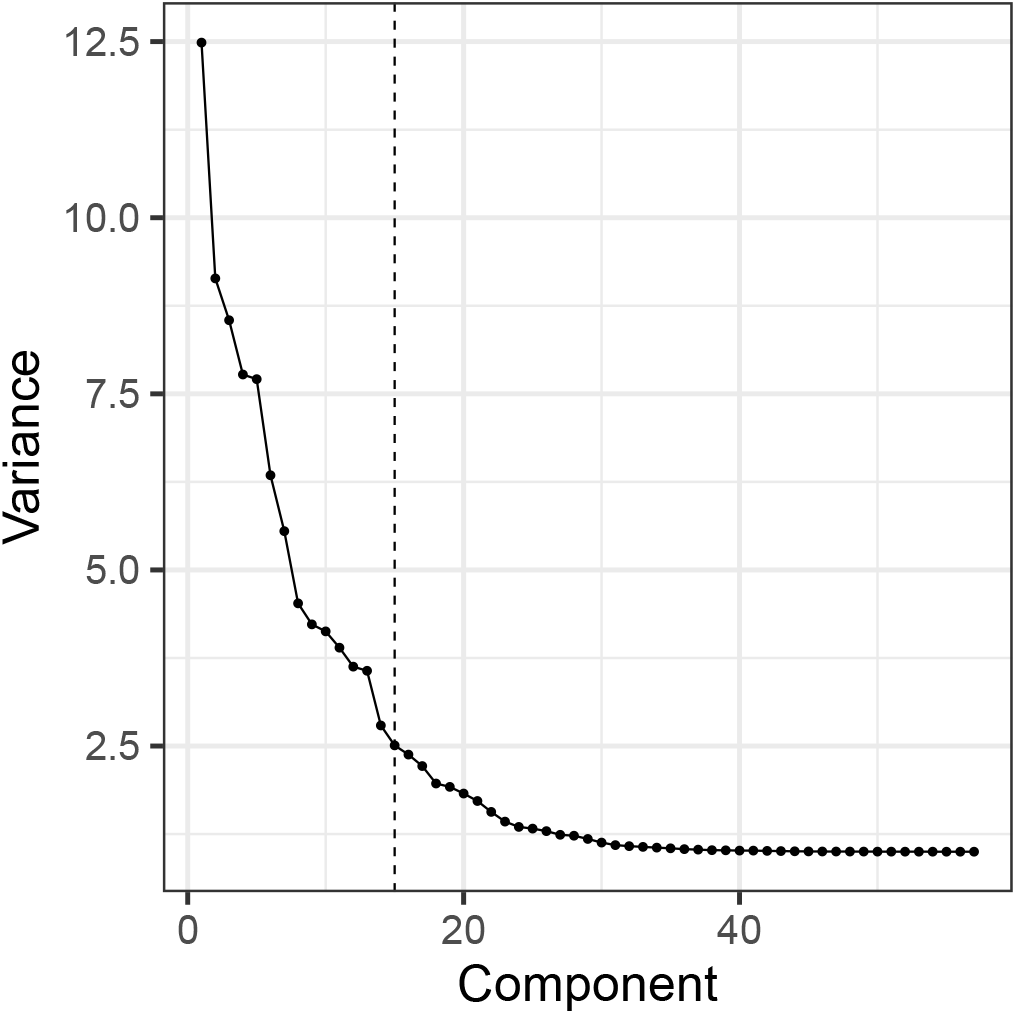
Variance of the single cell data along each orthogonal component of LDA. The held-out validation set is **not** included in the variance computation. The variance shows a large drop between the 13th and 14th LDA components. For this paper, we chose 15 LDA components for all downstream analyses.

For the quadratic program (2), we used the top 15 LDA components of the reference cohort as the reference matrix **X**. For the validation cohort, we first generate pseudobulk samples by summing gene counts across cells, then normalize the summed counts and successively apply the PCA and LDA rotation matrices to obtain ***µ***_*s*_.

**Figure A.2:**
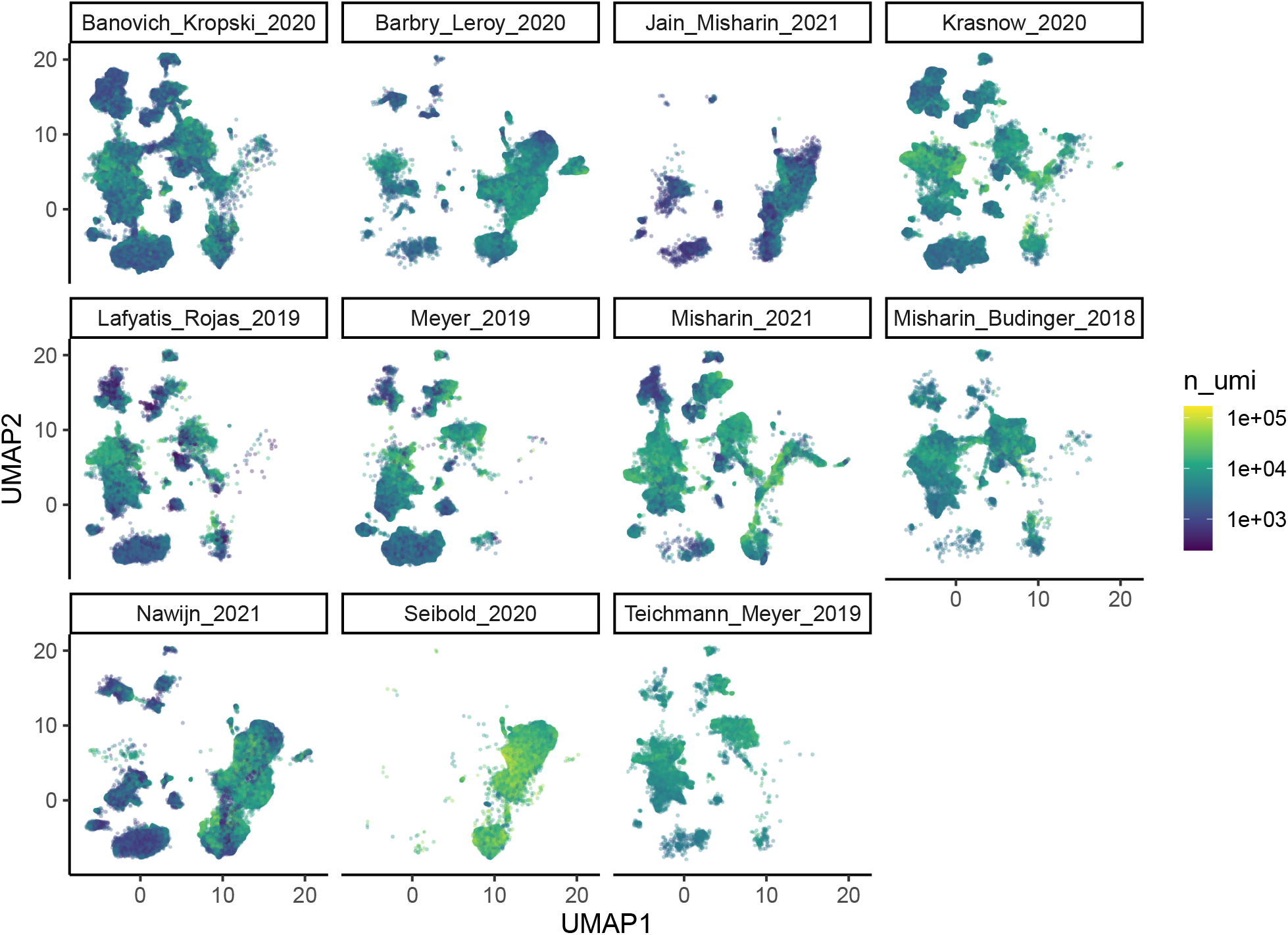
Number of UMIs per cell across the studies in the HLCA dataset. The studies have different sequencing depths and contain different cell types, which complicates estimating cell-specific size factors to convert between read-level and cell-level abundances in bulk deconvolution.

### A.2 Number of Alveolar macrophages vs Alveolar macrophage CCL3+ cells

At the finest resolution of celltype annotation, the bulk deconvolution results of Figure 1b were unable to distinguish the “Alveolar macrophage CCL3+” subpopulation from “Alveolar macrophages”. This was in part due to the rarity of “Alveolar macrophage CCL3+” in the reference set, which causes none of the top 15 LDA components to separate these populations well (Figure A.5). See Table A.2 for the number of cells from these subpopulations in each cohort.

**Table A.2:** Number of cells with celltype “Alveolar macrophages” or “Alveolar macrophage CCL3+”. Our bulk deconvolution results were unable to distinguish these 2 subpopulations (Figure 1b), assigning both cell types as “Alveolar macrophages”. This is in part due to the rarity of “Alveolar macrophage CCL3+” in the reference set (291 cells, or 0.4%), in comparison to the validation set (5785 cells, or 9.4%).

| Cohort | Alveolar macrophages | Alveolar macrophage CCL3+ |
| --- | --- | --- |
| Validation (Krasnow_2020) | 8597 (14%) | 5785 (9.4%) |
| Reference | 59890 (11.4%) | 2280 (0.4%) |

### A.3 Using Lagrangian dual solution for model diagnostics

In Figure A.4, we illustrate how the Lagrangian dual solution can be used to inspect the influence of each moment constraint. For example, the dual variable for the 14th Linear Discriminant is large, especially in the sample “distal 3”. That discriminant separates monocytes from macrophages (Figure A.5), which suggests that small perturbations of the bulk counts may cause the inferred weights to shift between these types of myeloid cells.

**Figure A.3:**
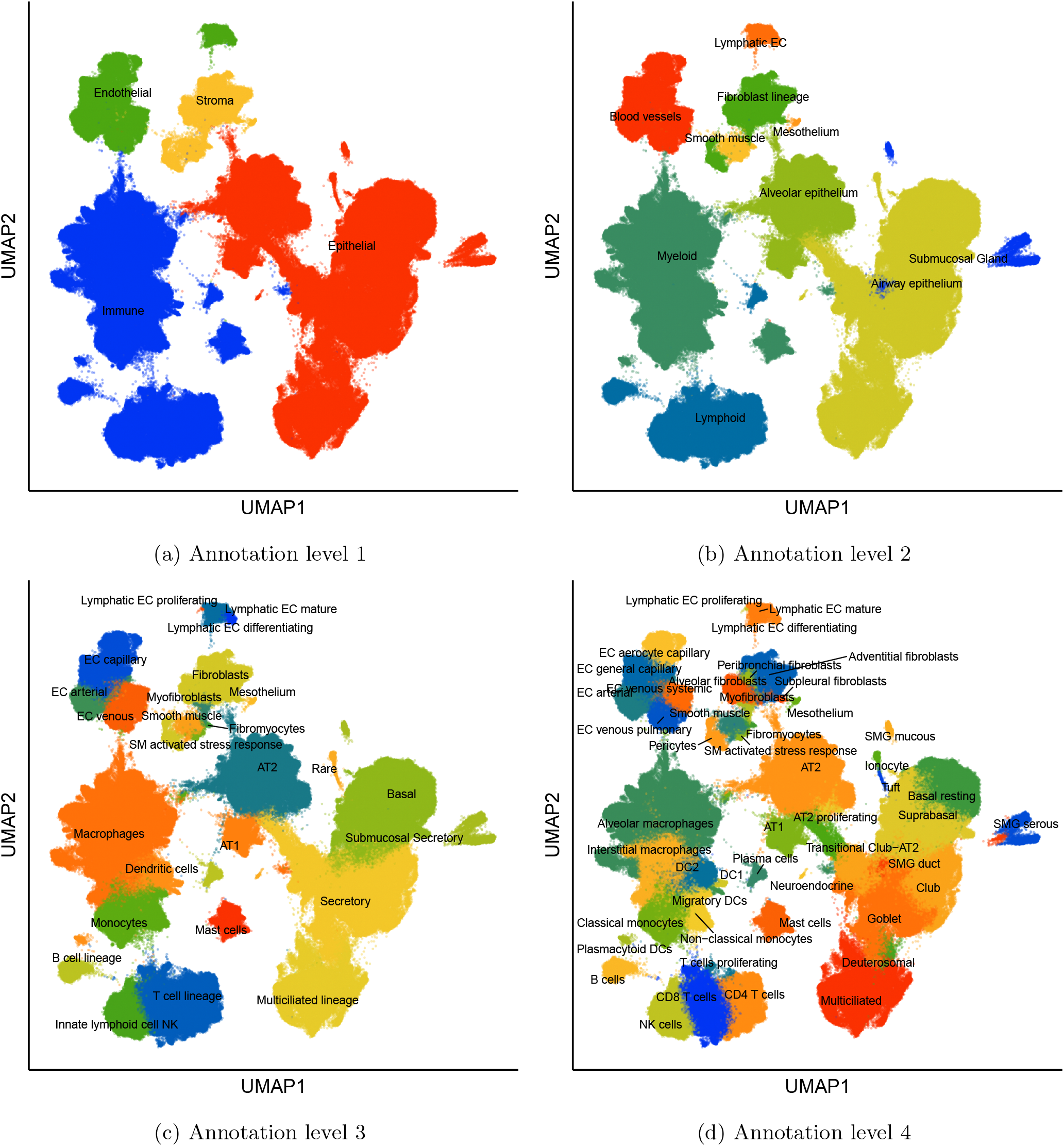
HLCA celltypes at annotation resolutions 1-4.

**Figure A.4:**
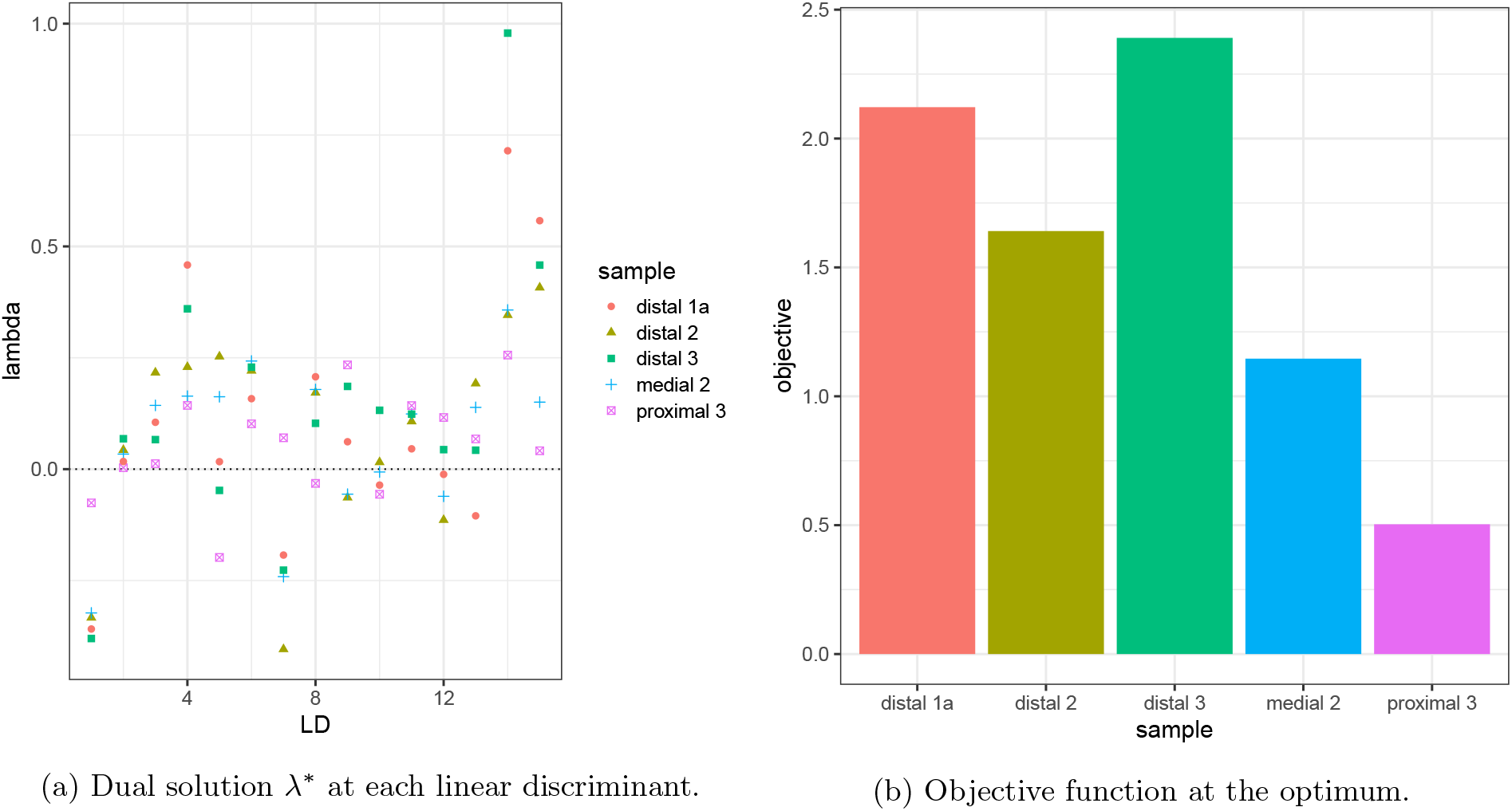
Dual variables and objective at the optimum for the bulk deconvolution example. The dual variables are the gradient of the objective and can be used to identify which features have a strong influence on the solution.

### A.4 Comparison of α values on deconvolution results

In Figure A.6, we compare deconvolution results on the HLCA example (using the downsampled 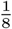 version of the reference, Table A.1) for *α*-divergences with *α* = 1, 1.5, 2. Note that *α* = 1 corresponds to Maximum Entropy, while *α* = 2 corresponds to the Pearson divergence. *α* = 2 yields the sparsest solution, while *α* = 1 yields the least sparse solution (and indeed, Maximum Entropy will never return probabilities of zero, see Jaynes (1957)). We also note that *α* = 1.5 was the most challenging to fit, both in runtime (taking 3 minutes, compared to about 5 seconds for Maximum Entropy and Pearson divergence), as well as in robustness – CVXPY’s default ECOS solver failed to converge, so we used the Clarabel solver (Goulart and Chen, 2024) instead (though note that CVXPY has announced it will be switching to Clarabel as the default solver in version 1.5).

**Figure A.5:**
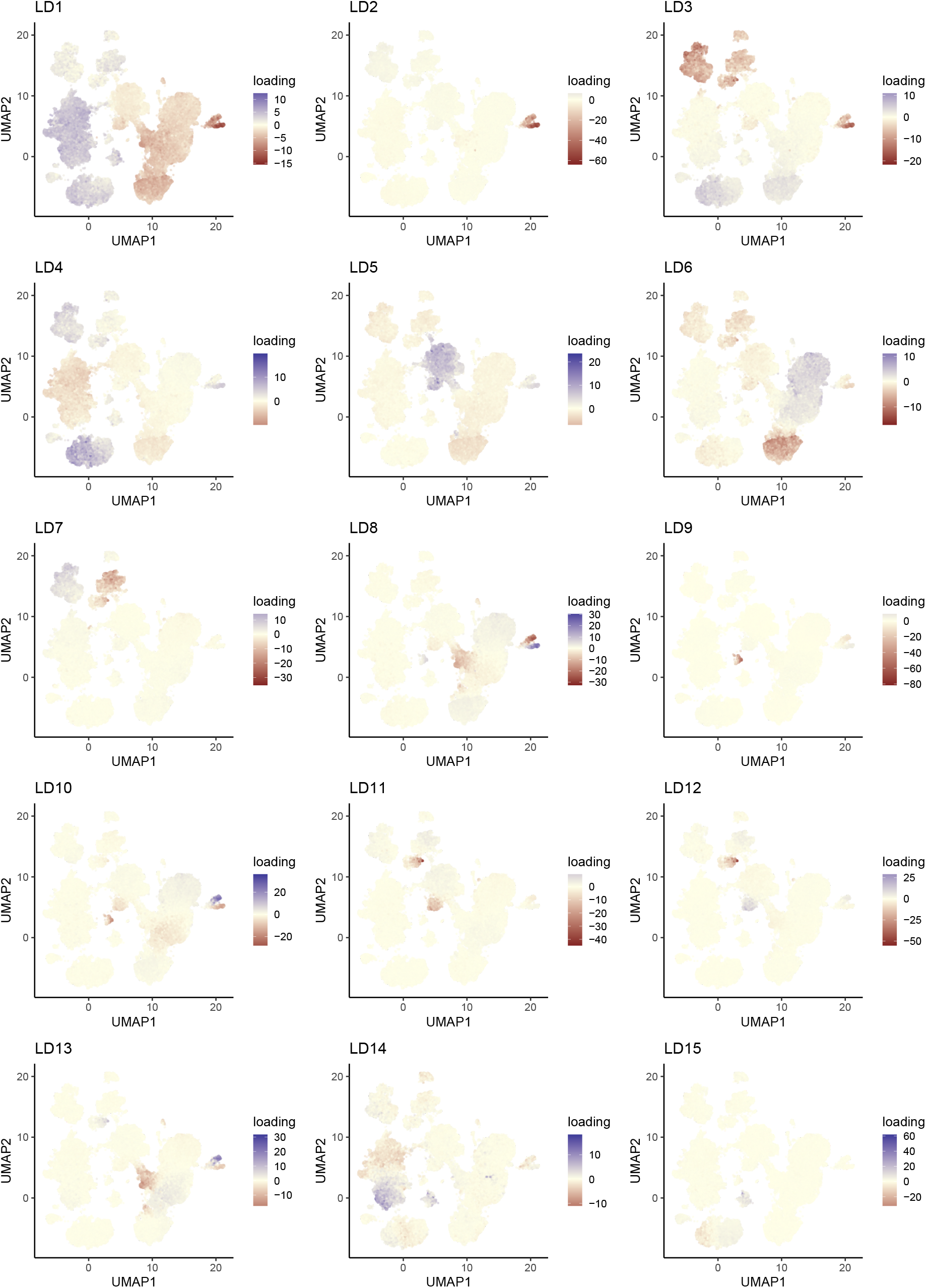
Cell loadings of each Linear Discriminant on the reference set.

**Figure A.6:**
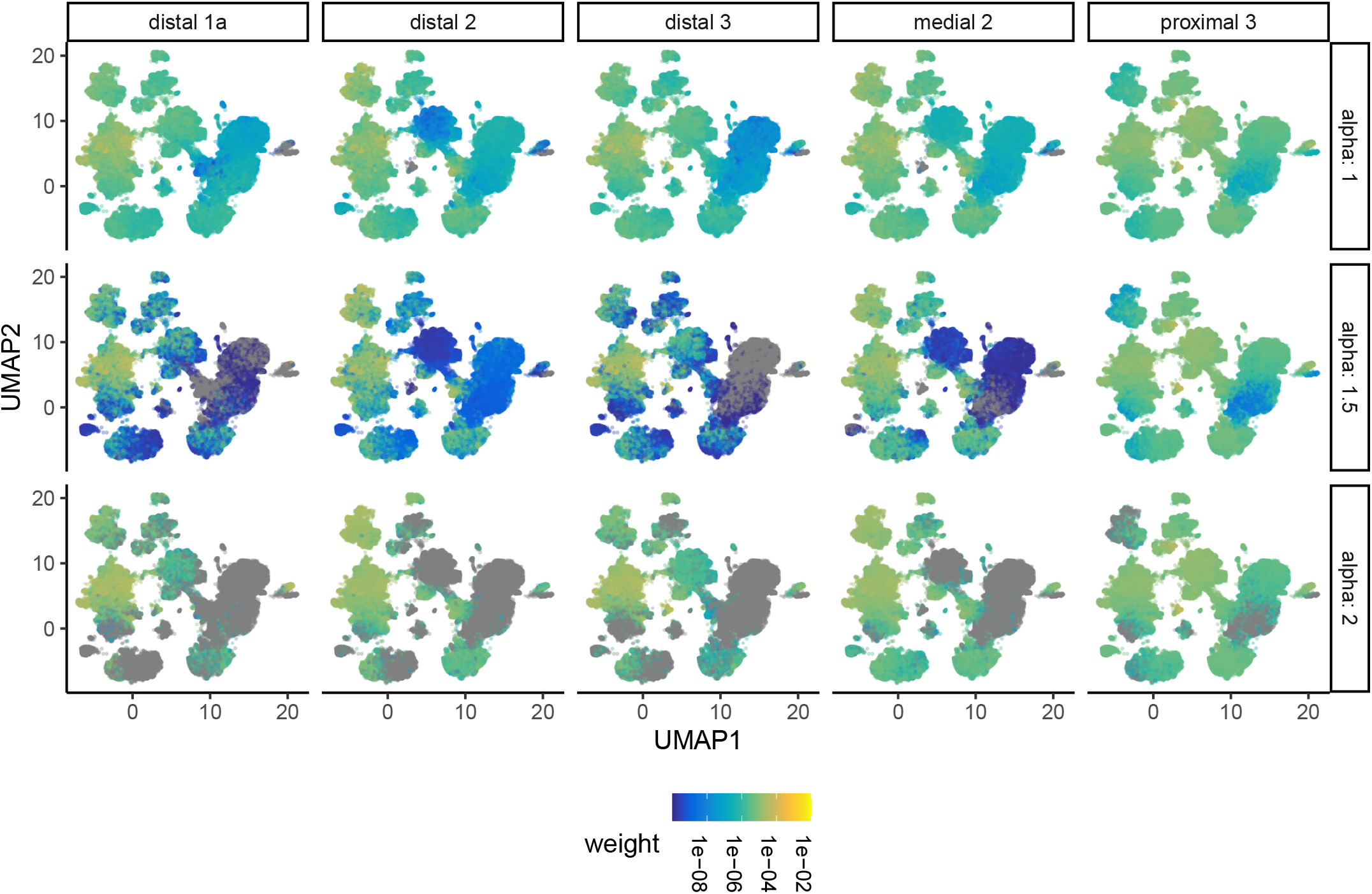
Bulk deconvolution results for the HLCA data for different values of *α*. For visualization purposes, weights below 1e-10 are set to 0, and appear as grey points.

